# Intraoperative electrical stimulation of the human dorsal spinal cord reveals a map of arm and hand muscle responses

**DOI:** 10.1101/2022.01.29.478182

**Authors:** James R. McIntosh, Evan. F. Joiner, Jacob L. Goldberg, Lynda M. Murray, Bushra Yasin, Anil Mendiratta, Steven C. Karceski, Earl Thuet, Oleg Modik, Evgeny Shelkov, Joseph M. Lombardi, Zeeshan M. Sardar, Ronald A. Lehman, Christopher Mandigo, K. Daniel Riew, Noam Y. Harel, Michael S. Virk, Jason B. Carmel

## Abstract

While epidural stimulation of the lumbar spinal cord has emerged as a powerful modality for recovery of movement, how it should be targeted to the cervical spinal cord to activate arm and hand muscles is not well-understood, particularly in humans. We sought to map muscle responses to posterior epidural cervical spinal cord stimulation in humans. We hypothesized that lateral stimulation over the dorsal root entry zone would be most effective, and responses would be strongest in the muscles innervated by the stimulated segment. Twenty-five people undergoing clinically indicated cervical spine surgery were consented to map motor responses. During surgery, stimulation was performed in midline and lateral positions at multiple exposed segments; six arm and three leg muscles were recorded on each side of the body. Across all segments and muscles tested, lateral stimulation produced stronger muscle responses than midline despite similar latency and shape of responses. Muscles innervated at a cervical segment had the largest responses from stimulation at that segment, but responses were also observed in muscles innervated at other cervical segments and in leg muscles. The cervical responses were clustered in rostral (C4-C6) and caudal (C7-T1) cervical segments. Strong responses to lateral stimulation are likely due to the proximity of stimulation to afferent axons. Small changes in response sizes to stimulation of adjacent cervical segments argues for local circuit integration, and distant muscle responses suggest activation of long propriospinal connections. This map can help guide cervical stimulation to improve arm and hand function.

**New and noteworthy:** A map of muscle responses to cervical epidural stimulation during clinically indicated surgery revealed strongest activation when stimulating laterally compared to midline, and differences to be weaker than expected across different segments. In contrast, waveform shapes and latencies were most similar when stimulating midline and laterally indicating activation of overlapping circuitry. Thus, a map of the cervical spinal cord reveals organization and may help guide stimulation to activate arm and hand muscles strongly and selectively.

**Graphical abstract:** 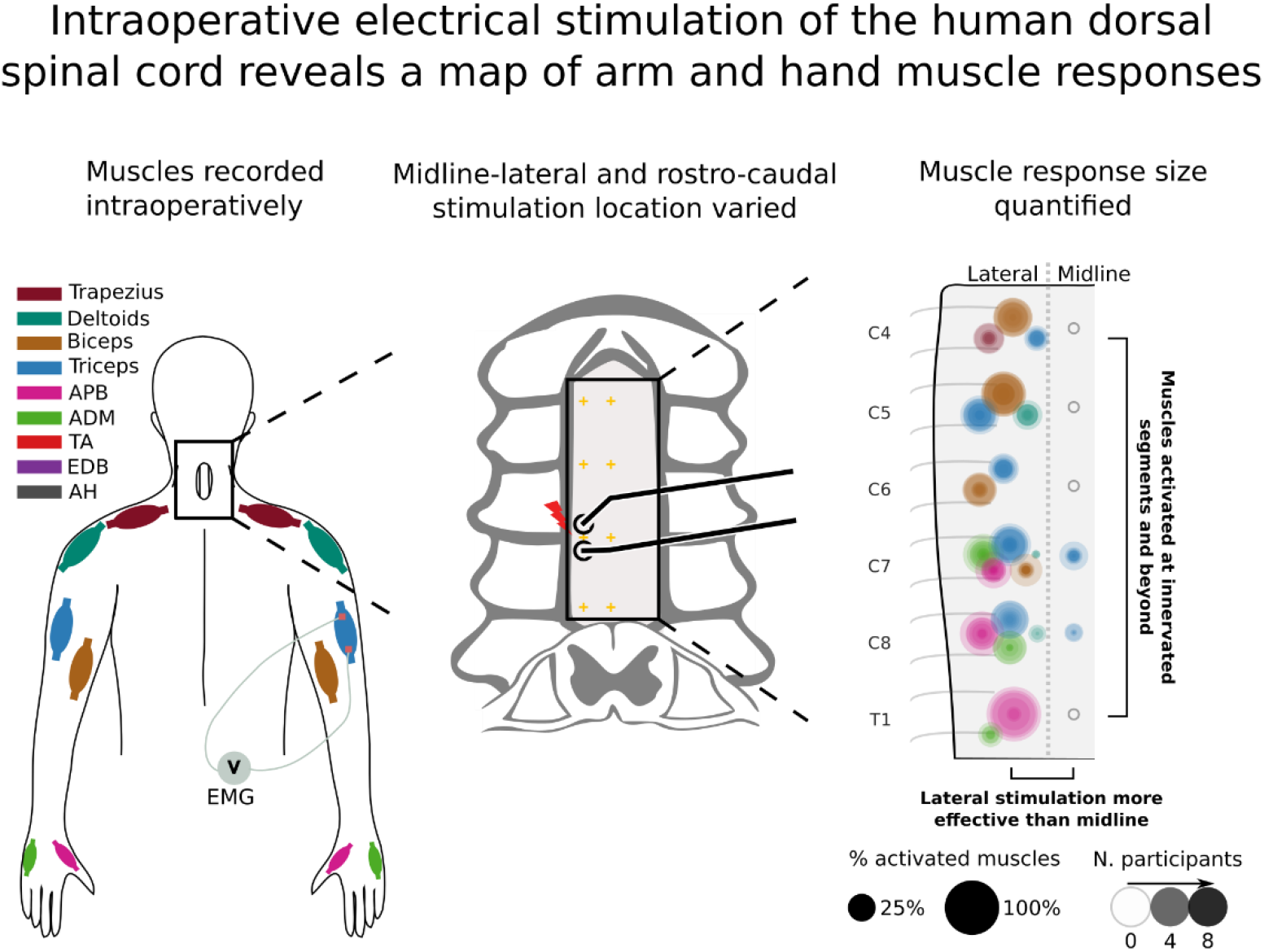

## 1 Introduction

Epidural stimulation has emerged as a way to modify spinal cord circuits for movement recovery (1–5). These studies largely targeted the lumbar spinal cord, with its relatively well-defined central pattern generator (6, 7). In contrast, the circuit-level logic of where and how to stimulate the cervical spinal cord is not as well known. Since hand function is the top priority of people with cervical spinal cord injury (SCI) (8, 9), interventions are under development to target the cervical spinal cord (10–15). Effective stimulation of the cervical spinal cord may be more difficult than for the lumbosacral spinal cord given its large behavioral repertoire and poorly understood intrinsic programs (16). The current study used electrical stimulation during clinically indicated cervical spine surgery to improve understanding of cervical spinal cord circuits.

Some of the insights learned from lumbar epidural stimulation apply to the cervical spinal cord. Stimulation of the dorsal spinal cord at low intensity evokes muscle responses via large-diameter afferents and not by direct activation of motoneurons (17, 18). Electrical stimulation of afferents activates motoneurons through circuits involved in reflexes (19, 20). This mechanism has been corroborated by mathematical modeling of current flow (21–23) as well as in inactivation studies (14). Consistent with this model, we have shown that stimulating near the dorsal root entry zone is more effective than stimulating at midline in the cervical spinal cord of rats (24).

Dorsal epidural stimulation of a cervical segment activates muscles innervated at that segment with spread to adjacent segments. In work performed mostly in monkeys, muscle responses to epidural stimulation of the cervical spinal cord (23, 25) broadly correspond to the distribution of motor pools throughout the cervical cord (26, 27). The spread of responses beyond the stimulated segment may be due to spread of afferents (28, 29) or variability in motoneuron to muscle connections (30). For example, with the exception of the separation between the biceps and triceps, Greiner et al. (23) observed a considerable overlap in the activated muscles at each cervical spinal segment in macaque monkeys. When stimulating dorsally in humans this breadth of activation of individual muscles across multiple segments is present; however, some additional divergence has also been found (31), with the most extreme observation being of leg muscle activation to dorsal epidural cervical stimulation (32). Leg muscle activation has also been observed in transcutaneous dorsal cervical stimulation (33).

These data led us to hypothesize that epidural stimulation of each cervical segment would activate large diameter afferents, causing strongest contraction of muscles innervated at that segment when stimulating laterally near the dorsal root entry zone. The detailed predicted results are shown in Fig. 1b, with larger circles representing larger MEPs provoked by stimulation at that location in biceps (brown) or triceps (blue). Similar to our studies in rats (24), we predicted that lateral stimulation of the spinal cord would be more effective than midline stimulation. The strongest responses would be observed at the segment of innervation (e.g. C5 and C6 for biceps (34) and C7 for triceps) with spread from there to adjacent segments. Taken together, we expected that there would be a larger change in the size of motor evoked potentials when comparing midline to lateral stimulation within each segment than when comparing rostral-caudal cervical segments. Despite these differences, we expected that midline and lateral stimulation would activate similar circuits, and we tested this by comparing MEP waveform shape and onset latency. The results are expected to help target cervical epidural stimulation to activate or modulate arm and hand muscles.

**Figure 1.**
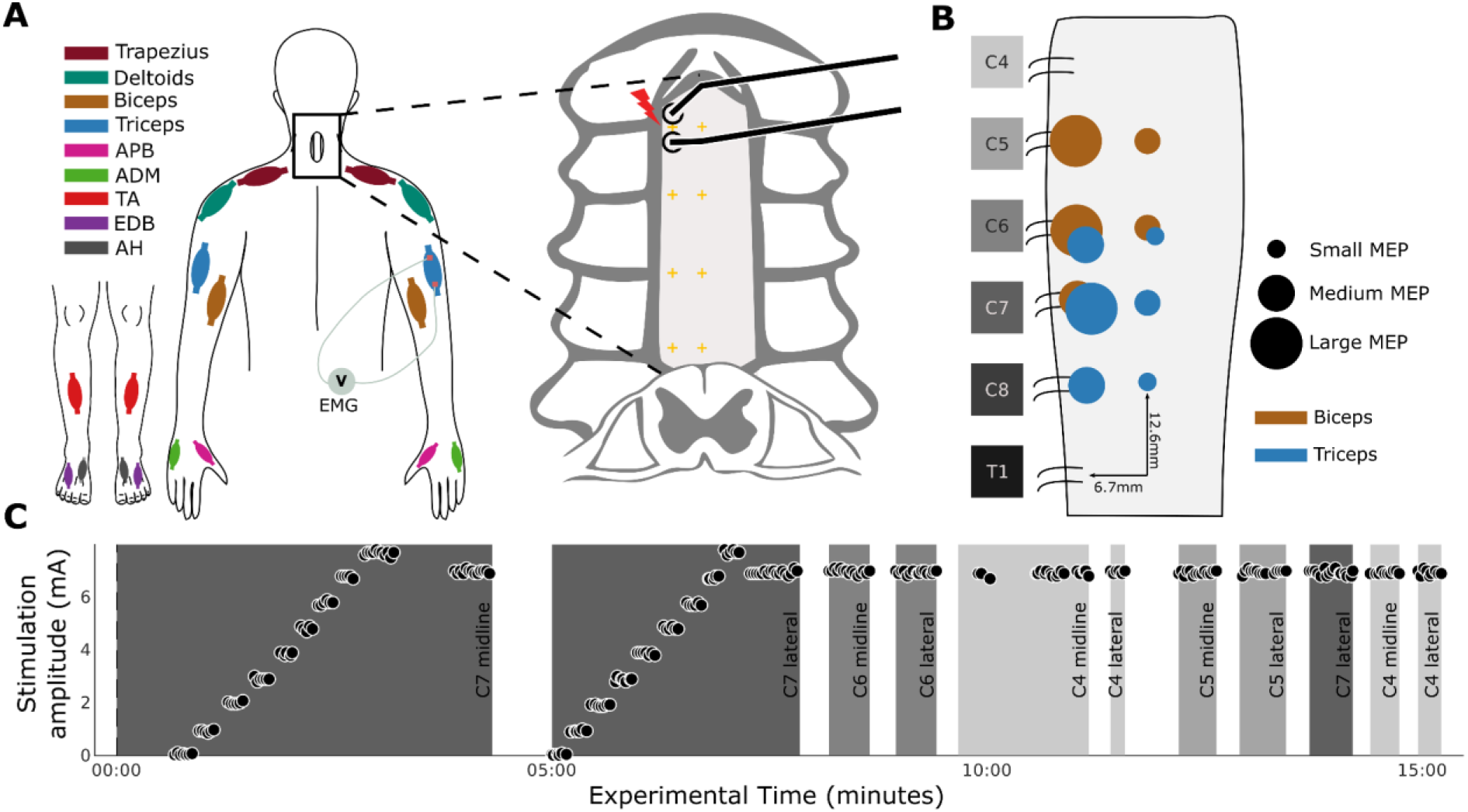
Epidural stimulation experiment during posterior cervical spine surgery. **(A)** Once the dura was exposed by laminectomy, a handheld or catheter electrode was used to stimulate at multiple segmental locations in either the lateral or midline position. Colors correspond to different recorded muscles (see legend). **(B)** Hypothesized responses to epidural stimulation of the cervical cord: 1) Lateral stimulation over the dorsal root entry zone was expected to be more effective than stimulation over the midline (large circles, representing high activation are lateral, while smaller circles are placed along the midline). 2) Muscle activation was expected to be relatively localized in the rostral caudal direction; however, some spread of this activation was expected. **(C)** Example sequence of the 15 minute experiment. Initially, the electrode was placed along the midline, over the dura at a segment aligned with the expected location of the root entry (C7 midline for this experiment). The stimulation intensity was ramped up with two purposes: 1) in search of the threshold of a target muscle. 2) To gather threshold information for multiple muscles in a single sweep. Once the threshold was found stimulation was performed at 1.2× that value The procedure was then repeated laterally to enable a midline-lateral comparison. Stimulation was performed at multiple segments at both midline and lateral locations.

## 2 Materials and methods

### 2.1 Experimental design

Among people undergoing clinically indicated spine surgery, epidural electrical stimulation of the exposed segments of the cervical spinal cord was performed, and motor evoked potentials (MEPs) from arm and leg muscles were recorded (Fig. 1A). Surgical time was not extended by the experimental procedure for longer than 15 minutes to limit the surgical and anesthetic risk of increased intraoperative time (35). To test whether lateral stimulation would be more effective than midline, stimulation at each of these sites was compared (Fig. 1B). To test whether stimulation at one cervical segment would produce the largest responses in muscles innervated at that segment, stimulation at multiple cervical roots was performed. To test whether circuits activated at midline and lateral stimulation are more similar to each other than circuits activated at different rostral-caudal segments, we compared MEP onset latencies and correlations in waveform shape. To test whether moving the stimulating electrodes in the midline-lateral direction would produce larger changes than in the rostro-caudal direction, the size of MEP change over change in distance was computed for each direction (see Methods section “Comparison of responses at different cervical segments”). A representative experiment is shown in Fig. 1C. The primary outcome was the size of the MEP, and the secondary outcomes were the threshold and slope of the recruitment curve with increasing stimulus intensity, as well as the onset latency and shape of the resultant MEPs.

### 2.2 Confirmation of electrode position over the dorsal root entry zone

When placed laterally, electrodes were positioned straddling the expected location of the dorsal root entry zone in a rostral-caudal orientation as identified by bony and neural anatomical landmarks. Specifically, the exiting cervical nerve was identified at each relevant level as it entered the neural foramen just caudal to the pedicle. The electrodes were then placed at the site the dorsal root enters the lateral spinal cord. Electrode placement based on anatomical landmarks was subsequently confirmed to be over the dorsal root entry zone via image reconstruction and coregistration between preoperative MRI and intraoperative CT scans performed in one participant. Clinically indicated intraoperative-CT (Airo, Stryker Inc.) and pre-operative MRI (T2-weighted, Siemens) were gathered for this participant. During the course of the experiment, the location of the C7 root entry was inferred based on the bony landmarks of the spine. After placing the electrode at that site an intraoperative photograph was taken (Fig. 2A1). The location of the electrode was then translated to a location in the CT based on instrumentation inserted during surgery (Fig. 2A2). The CT and MRI images were then co-registered by performing a non-linear registration with the “Curvature Correction” software (Brainlab AG) to account for the different curvature of the spine in prone (operative) and supine (preoperative) participant positions (Fig. 2A3). The transformation generated in the registration was then also applied to the location of the electrode in the CT and shown in the MRI (Fig. 2A4).

**Figure 2.**
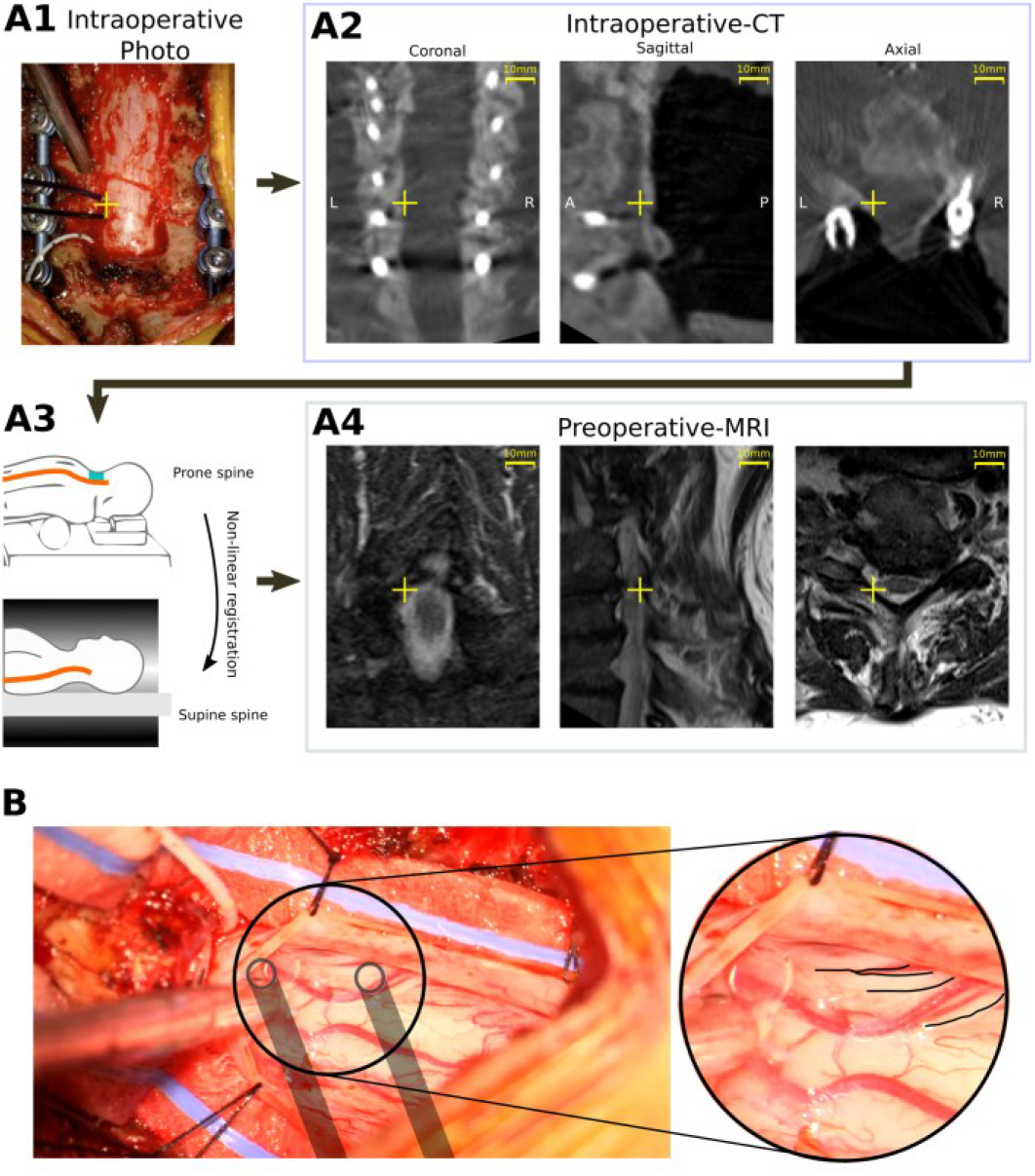
Validation of stimulation location relative to dorsal root entry zones. **(A)** Transformation of electrode stimulation site from physical location to MRI space. (A1) A photograph was used to identify the position of electrodes relative to instrumentation. (A2) This location was translated to a CT scan used to confirm implant location (yellow crosshatch). (A3) CT and MRI images were co-registered by performing a non-linear segmental registration (Brainlab AG) to account for differences in spinal alignment found in prone (operative) and supine (preoperative) positions. (A4) Electrode position in MRI near targeted C7 root verified location. **(B)** Exposed subdural region with overlaid location of stimulated location and dorsal roots highlighted in inset. Placement based on anatomical landmark identification was confirmed to position the electrodes over the dorsal root entry zones.

Confirmation of electrode position was also made visually in an additional experimental condition where stimulation was applied at the T1 root entry, below the surface of the dura and at matched epidural locations (Fig. 2B).

### 2.3 Participants

Participants were adult patients with cervical spondylotic myelopathy, multilevel foraminal stenosis, or intradural tumor requiring surgical treatment (Table 1, Table S1). Patients were enrolled from the clinical practices of the spine surgeons participating in the study. The study protocol was reviewed and approved by the institutional review boards of the two study sites, Weill Cornell Medical Center and Columbia University Irving Medical Center (ClinicalTrials.gov number: NCT05163639, participants were recruited after registration). Patients with stimulation devices in the neck or chest (e.g. vagal nerve stimulation, cardiac patients with pacemakers) or head and neck implants were excluded. Informed consent for participation in the study was obtained prior to surgery for every participant. Participants underwent standard of care preoperative clinical assessments. The modified Japanese Orthopaedic Association (mJOA) scores were used to assess the severity of myelopathy. These experiments were powered with effect size based on comparisons of midline and lateral stimulation in the rat (24). Conservatively assuming a reduced effect size by 50% compared to the rat experiments (Cohen’s *d* = 0.97 versus 1.94), the analysis indicated that to achieve 90% power using a Wilcoxon signed-rank test (*α* = 0.05) would require 14 participants. Additional participants were recruited to map responses from each cervical segment, with a minimum of 5 participants from each level of the cervical enlargement (C5-T1).

**Table 1.** Clinical characteristics of participants. See Supplemental Table 1 for associated experiment parameters.

|  | Age*/Sex | mJOA | Symptom onset | Dominant Syndrome | Unsteady Gait | Pain | Numbness | Weakness | Hyperreflexia |  | Sphincter dysfunction | T2 signal change |
| --- | --- | --- | --- | --- | --- | --- | --- | --- | --- | --- | --- | --- |
|  |  |  |  |  |  |  |  |  | Upper Extremities | Lower Extremities |  |  |
| P01 | 67M | 14 | Chronic | Myeloradiculopathy | No | Radiating | Yes | Yes | No | No | No | None |
| P02 | 48F | 14 | Chronic | Radiculopathy | No | None | Yes | Yes | Yes | No | No | C2-3, C4-5, C5-6 |
| P03 | 58M | 10 | Subacute | Myelopathy | Yes | None | Yes | Yes | No | No | Yes | C4-5, C5-6 |
| P04 | 54M | 17 | Chronic | Radiculopathy | No | Radiating | Yes | No | No | No | No | C3-4, C6-7 |
| P05 | 74F | 12 | Chronic | Myeloradiculopathy | Yes | Radiating | Yes | Yes | Yes | Yes | No | C3-7 |
| P06 | 71F | 11 | Chronic | Myelopathy | No | Radiating | Yes | Yes | Yes | No | No | C4-5, C5-6 |
| P07 | 62M | 10 | Chronic | Myelopathy | Yes | None | Yes | Yes | Yes | Yes | No | C6-7 |
| P08 | 83F | 14 | Chronic | Myeloradiculopathy | No | Radiating | No | Yes | Yes | Yes | No | C3-4, C4-5, C5-6 |
| P09 | 72F | 12 | Subacute | Myelopathy | No | Axial Neck | Yes | Yes | No | No | No | C3-4 |
| P10 | 70F | 10 | Chronic | Myelopathy | No | Radiating | Yes | Yes | Yes | No | No | C3-4, C4-5, C6-7 |
| P11 | 74F | 13 | Chronic | Myeloradiculopathy | No | Radiating | Yes | Yes | No | No | No | C3-4 |
| P12 | 50F | 11 | Chronic | Myeloradiculopathy | No | Radiating | Yes | Yes | Yes | No | No | C5-7 |
| P13 | 60M | 17 | Chronic | Radiculopathy | No | None | Yes | No | No | No | No | C3-4 |
| P14 | 74M | 15 | Chronic | Myelopathy | Yes | Axial neck | No | No | Yes | Yes | No | C4-5 |
| P15 | 75M | 16 | Chronic | Myeloradiculopathy | Yes | Axial neck | Yes | Yes | No | No | No | None |
| P16 | 54M | 17 | Chronic | Radiculopathy | No | Radiating | Yes | No | No | No | No | C6-7 |
| P17 | 71F | 11 | Chronic | Myeloradiculopathy | Yes | Axial neck and radiating | No | Yes | Yes | Yes | No | C3-4 |
| P18 | 56M | 11 | Chronic | Myeloradiculopathy | Yes | Axial neck and radiating | No | Yes | Yes | Yes | No | C2-3, C3-4, C5-6 |
| P19 | 39M | 10 | Chronic | Myelopathy | Yes | None | Yes | Yes | No | No | Yes | C4-5 |
| P20 | 78F | 14 | Chronic | Myelopathy | Yes | Axial neck and upper back | No | Yes | No | No | No | None |
| P21 | 72F | 12 | Chronic | Myelopathy | Yes | Axial neck | No | Yes | No | No | No | C3-4, C4-5, C6-7 |
| P22 | 76F | 10 | Chronic | Myeloradiculopathy | Yes | Axial neck and radiating | Yes | Yes | No | No | Yes | C5-6 |
| P23 | 49F | 17 | Chronic | Myeloradiculopathy | Yes | Axial neck and radiating | Yes | Yes | Yes | Yes | No | T1-2 |
| P24 | 79M | 11 | Chronic | Myelopathy | Yes | None | Yes | Yes | No | Yes | No | C4-5, C5-6 |
| P25 | 71M | 13 | Subacute | Myelopathy | Yes | None | Yes | Yes | No | No | No | C3-4, C4-5, C5-6 |

\*At the time of surgery. mJOA, modified Japanese Orthopaedic Association (mJOA) score.

### 2.4 Electrophysiology

Stimulation was performed with epidural electrodes in participants undergoing clinically indicated surgery, and the muscles recorded were those chosen based on the standard montage for cervical spine surgeries performed at our institutions. After anesthesia induction, only total intravenous anesthesia was used. No anesthetic adjustments were made during the 15 minutes dedicated to the experiment. Recording and stimulation were performed with Cadwell Elite/Pro, Cadwell IOMAX (Cadwell Inc.) or XLTEK Protektor32 (Natus Medical Inc.) intraoperative monitoring systems. The stimulation device chosen for a particular experiment was dependent on the study site and availability (see Table S1). The experimental procedure began once the dura was exposed and the epidural stimulation electrode was placed. Muscles were chosen for electromyogram (EMG) per standard of care (Fig. 1A). MEP responses were recorded with subdermal needles at a sampling rate between 6kHz and 10.4kHz and bandpass filtered between 10Hz and 2000Hz. Epidural spinal cord stimulation was performed in trains of 3 pulses with a hand-held double ball tip probe (2.3mm diameter contacts, 10mm spacing); single pulse and catheter electrode stimulation were also used at specific cervical segments (see Table S1). Stimulation with a 3-pulse train was used to reduce the intensity necessary to evoke an MEP in order to reduce current spread (36). In each case, the electrodes were oriented in the rostro-caudal direction (Fig. 1A, inset) with the cathode caudal. At each testing site, stimulation was delivered every 2 seconds, a frequency that we determined does not alter responses with repeated stimulation (data not shown).

#### 2.4.1 Comparison of midline versus lateral stimulation

Within each cervical segment, midline stimulation was compared with lateral stimulation. Midline electrode placement was determined by visual estimation and confirmed with measurement from the bony landmarks on either side. Electrodes were allowed to dimple the surface of the dura to approximately one-half the depth of the ball tip. Prior to the start of the experiment, we designated the lateral stimulation to be performed on the left or right to match each participant’s less impaired side based on clinical signs, symptoms and MRI; this was done to minimize interaction of neurological deficits on electrophysiological responses.

*Stimulation sites and intensity.* An example of one experiment is shown in Fig. 1C. To begin each experiment, the stimulation electrode was placed on midline at the most caudal segment exposed during surgery. Stimulation intensity was incrementally increased from 0 to 8mA in order to assess the activation threshold and estimate the subsequent recruitment curve (minimum 10s per stimulation intensity; first panel Fig. 1C). Threshold was defined as the lowest stimulation intensity to produce an MEP in the most responsive muscle at the initially tested segment. The threshold of this muscle (Table S1) was used to set the fixed stimulus intensity (120% of threshold) for 30 seconds of stimulation at other segments. The experiments proceeded by repeating the stimulation intensity ramp and fixed intensity stimulation at the equivalent lateral site (second panel of Fig. 1C).

#### 2.4.2 Comparison of responses at different cervical segments

MEPs at multiple segments were compared using the fixed intensity, which was repeated at more rostral segments (third panel onwards of Fig. 1C). This stimulation protocol was applied in all experiments. Additional experimental conditions were tested (see supplemental Table S1) using single pulse stimulation and a flexible catheter electrode (1.3mm contacts, 15mm spacing, Ad-Tech Medical Instrument Corp). The catheter could be inserted below the lamina to allow access to segments not directly exposed surgically. We confirmed that the data from catheter stimulation was similar to the data recorded with ball electrodes and that exclusion of this data would not substantially change the results (data not shown).

### 2.5 Data analysis

Data was exported from proprietary intraoperative monitoring software to MATLAB (R2020b) and Python (v3.8) where analysis was performed. MEPs were quantified using the rectified area under the curve (AUC) calculated in a window between 6.5ms and 75ms after the start of the first stimulation pulse. In order to indicate the absence or presence of MEPs (dashed line in Fig. 8, cutoff in Fig. Fig. 10) an equivalent estimate of the 50μV threshold was used as is typical in non-invasive studies (37). This value was estimated by regressing AUC onto the peak-peak MEP size, resulting in an AUC of 0.33μVs.

#### 2.5.1 Statistical analysis

Values are reported as mean ±2×SEM except in cases where the median is used. Non-parametric statistical tests are used throughout (Wilcoxon rank-sum and signed-rank tests, alpha = 0.05).

#### 2.5.2 Artifact rejection

Rejection was based on principal component analysis and confirmation by a human observer The principal components were computed for a specific muscle and electrode location across multiple stimulation intensities. These principal components captured the shape of the MEPs, and were regressed with each MEP. The root-mean-square of the regression error was then used to rank the responses. This ranking sorted the responses so that the most dissimilar waveform shapes were ranked highest. A manually adjusted sliding scale was then used to reject the highest ranked traces that did not appear physiological under visual inspection: deflections in baseline, spread of stimulation artifact into the evoked response, excessive line noise and fluctuations that were not time-locked to other responses. This led to 1,728 of the 112,989 MEPs (1.5%) being rejected.

#### 2.5.3 Midline-lateral comparisons

The stimulation efficacy of midline stimulation was compared to lateral stimulation in 4 ways. First, the MEPs for midline and lateral stimulation at 120% of midline threshold were compared for each individual participant (Fig. 3, insets), using the Wilcoxon rank-sum test, Bonferroni corrected for the number of participants (n = 14, all participants with triple pulse stimulation at midline and lateral locations). Second, the mean AUC for midline and lateral stimulation across the 14 participants was compared using the Wilcoxon signed-rank test (Fig. 3A). The same method applied to the triceps muscle MEPs (Fig. 3) was applied to the tibialis anterior responses in Fig. 5. Third, to ensure that differences observed in specific muscles at specific segments of stimulation were general trends, all of the responses of the 6 arm and hand muscles were plotted in Fig. 4. The percentage of responses that were larger for lateral stimulation was computed (Fig. 4, inset for each muscle). The fourth comparison of midline versus lateral stimulation was performed using recruitment curves. The curves were used to estimate thresholds for MEPs and the rate of change of MEPs with stimulation intensity (slope). A function that approximates a linear recruitment curve was used (see section 2.5.7). In a single participant where epidural and subdural stimulation sites were tested, midline and lateral conditions were compared using recruitment curve thresholds (Fig. 7).

**Figure 3.**
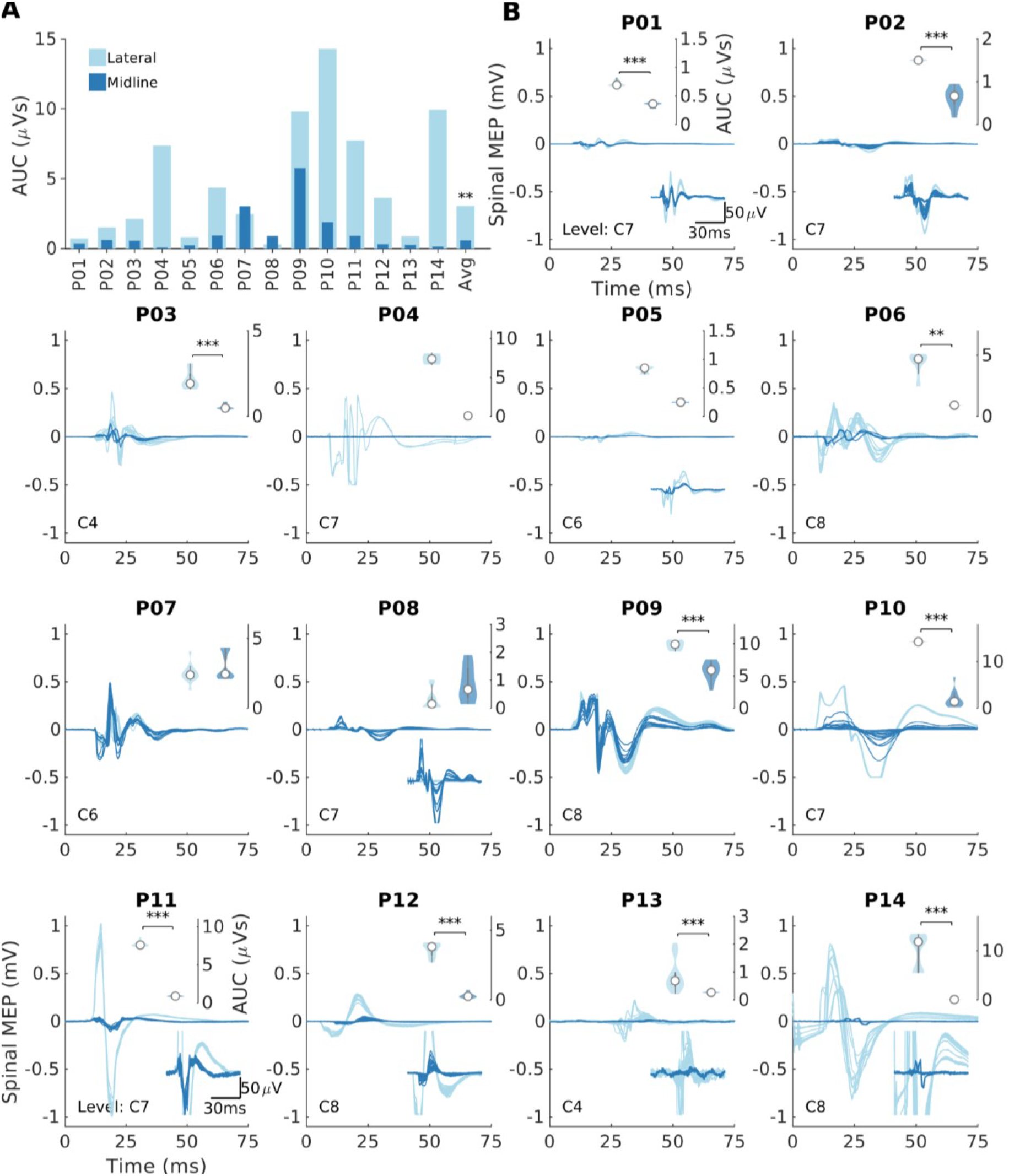
Lateral stimulation is more effective than midline stimulation. **(A)** Summary of the rectified area under the curve (AUC) of the triceps muscle at the most caudal segment where a response was present. Midline simulation (dark color) and lateral (light color) stimulation were performed at the fixed intensity for each participant. Lateral stimulation produced consistently larger responses for the most caudal responsive segments. A signed-rank test was applied between the midline and lateral conditions. Average bar represents the median. **(B)** Individual MEPs that compose the summary plot for individual participants. Bottom inset of each panel represents a magnification of the MEP in cases where the response is small. Top inset of each panel shows a violin plot of AUC (white circle represents the median). Within-participant Wilcoxon rank-sum tests were conducted in individual participants (\**p* < 0.05, \*\**p* < 0.01, \*\*\**p* < 0.001, Bonferroni corrected for multiple comparisons). Note that for P04, P09 and P10 the signal saturated at 0.5mV due to a limitation of the recording hardware.

**Figure 4.**
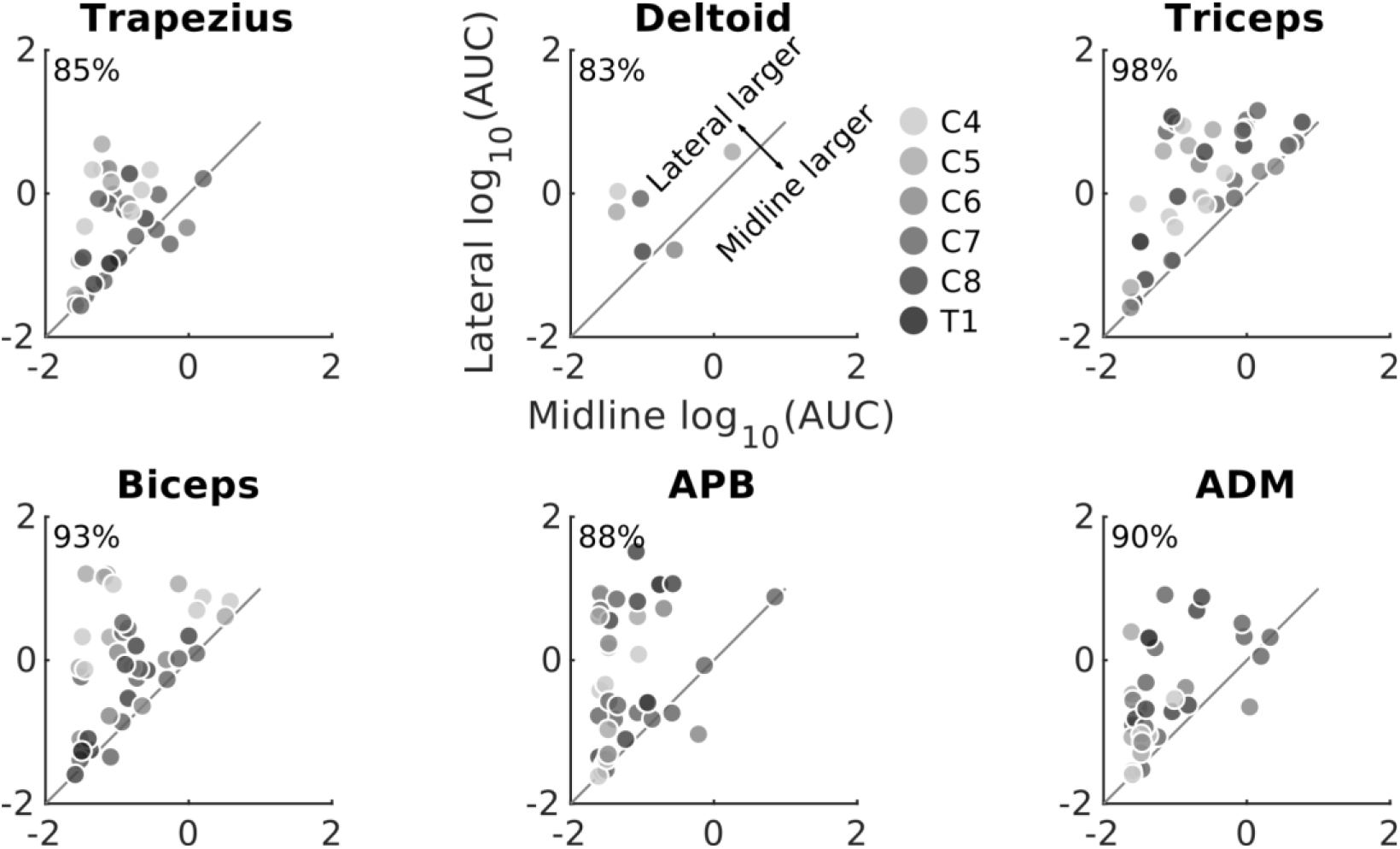
Lateral stimulation is more effective than midline stimulation regardless of stimulation segment or recorded muscle. AUC from all segments and muscles for all participants in whom both midline and lateral stimulation was performed. AUC was larger when stimulation was applied laterally than at midline in the majority of tested cases. The percentage of MEPs that were larger in lateral than midline stimulation is shown in the upper left corner of each plot.

**Figure 5.**
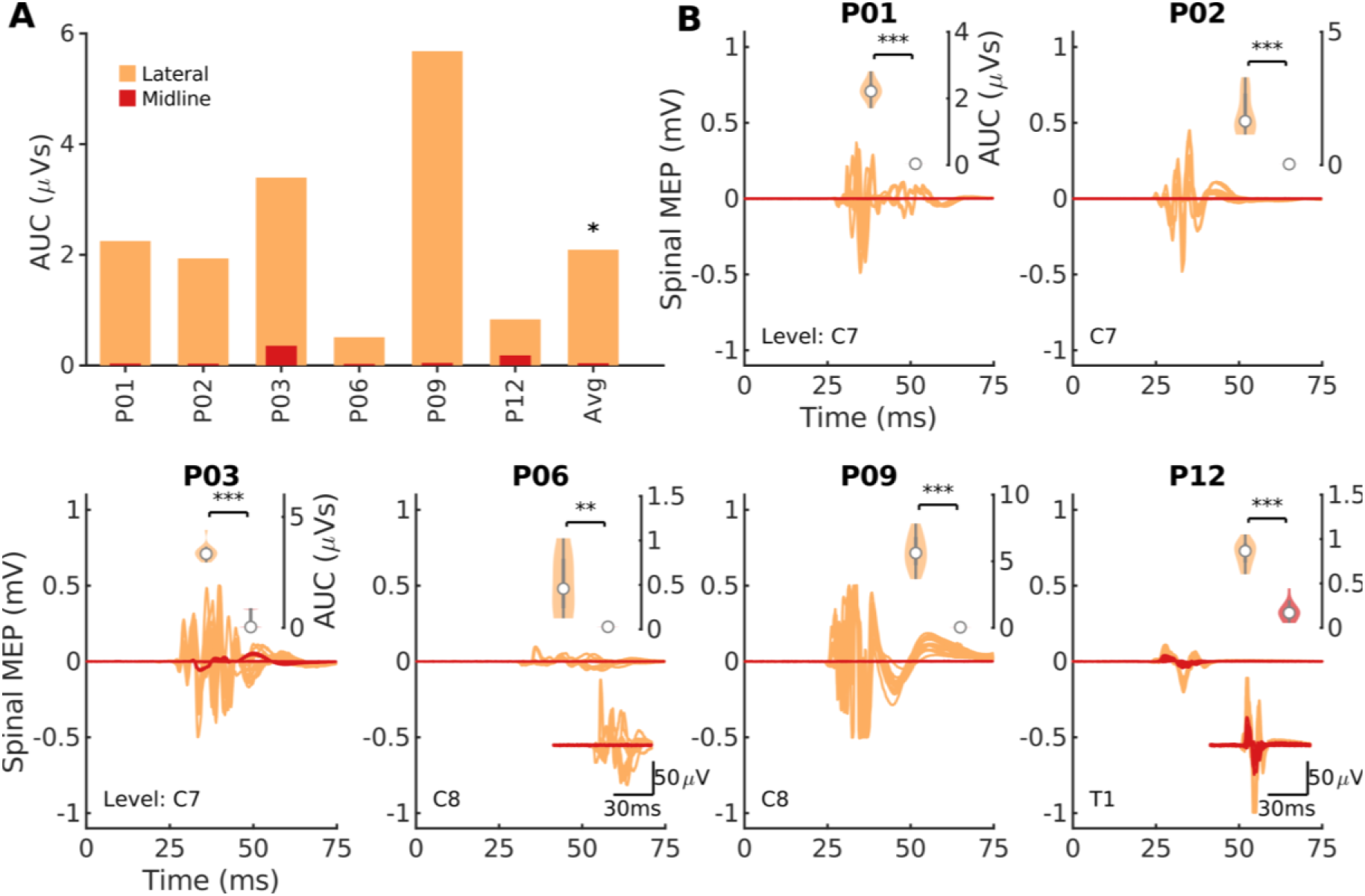
A subset of participants display activation in their leg muscles when stimulation is applied in the cervical cord. **(A)** Summary of the rectified area under the curve (AUC) of the tibialis anterior (TA) muscle at the most caudal segment where a response was present. Midline simulation (dark color, red) and lateral (light color, orange) stimulation was performed at the fixed intensity for each participant. Motor evoked potentials (MEPs) are shown for the most caudal responsive segment. In a subset of participants that display activation of tibialis anterior (AUC greater than 0.33μVs), lateral stimulation produced consistently larger responses. A signed-rank test was applied between the midline and lateral conditions; the average bar represents the median value. **(B)** Individual spinal MEPs that compose the summary plot for individual participants. Bottom inset of each panel represents a magnification of the MEP in cases where the response was small. Top inset of each panel shows a violin plot of AUC (white circle represents the median). Within participant Wilcoxon rank-sum tests were conducted in individual participants (\**p* < 0.05, \*\**p* < 0.01, \*\*\**p* < 0.001, Bonferroni corrected for multiple comparisons). Note that for P09 the signal saturated at 0.5mV due to a limitation of the recording hardware.

**Figure 6.**
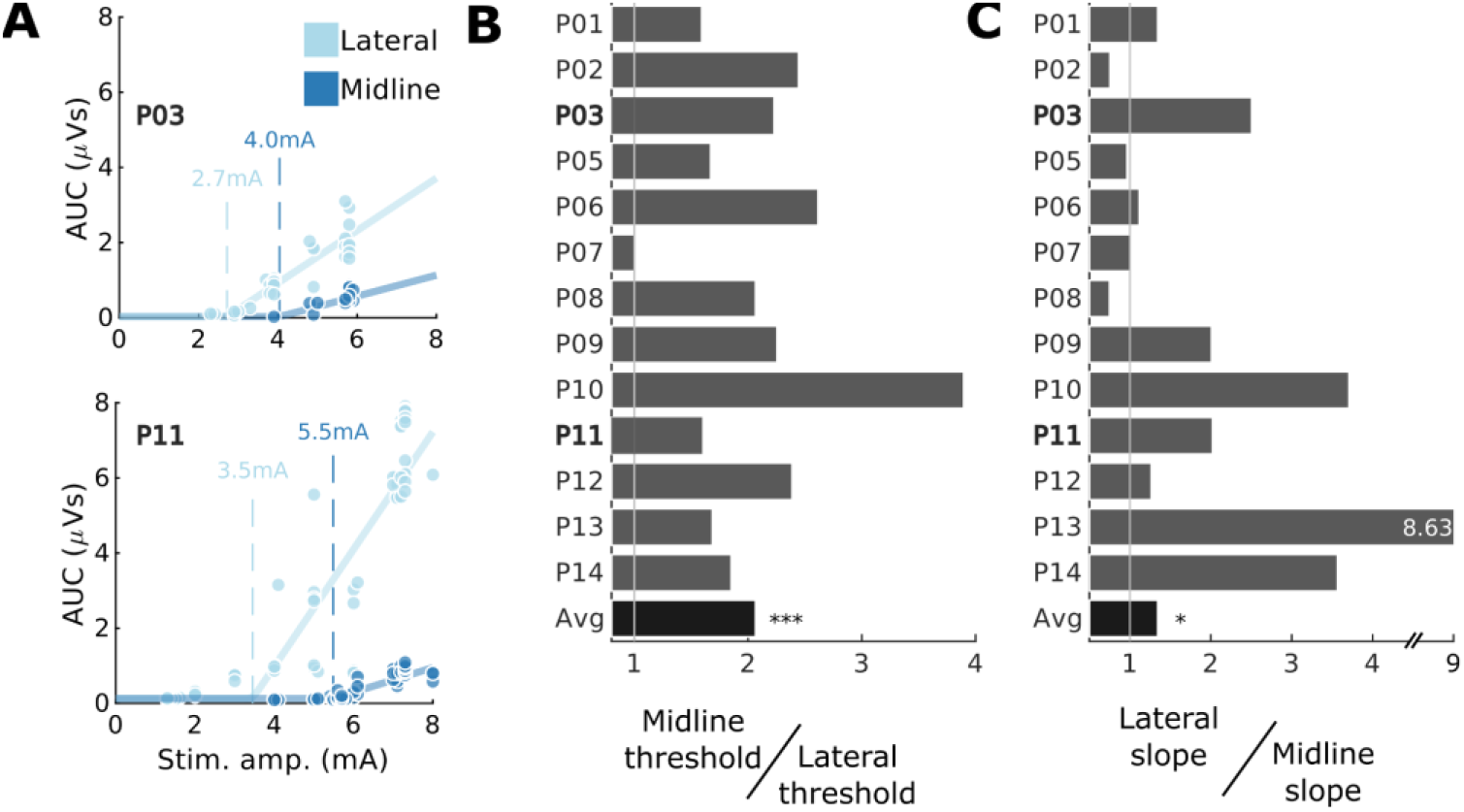
Larger lateral MEPs are driven by both a reduction in threshold and increase in slope. **(A)** The threshold of muscle activation was determined by fitting functions to sections of data where multiple stimulation intensities were used. Circles indicate individual data points (light color, lateral; dark color, midline), solid lines show example fitted functions, and dashed lines show estimates of threshold. Triceps muscle is being shown for two participants as indicated in inset text. **(B)** The relative efficacy of lateral stimulation can be summarized as midline threshold ÷ lateral threshold (lower lateral threshold indicates higher lateral efficacy). The majority of participants showed lower lateral than midline threshold. **(C)** Lateral to midline slope ratio (lateral slope ÷ midline slope). The majority of participants showed steeper recruitment slopes from stimulation at lateral sites. Note that one participant has been omitted (P04) as there was insufficient range in tested stimulation intensities to perform model fitting. Average bar represents the median.

**Figure 7.**
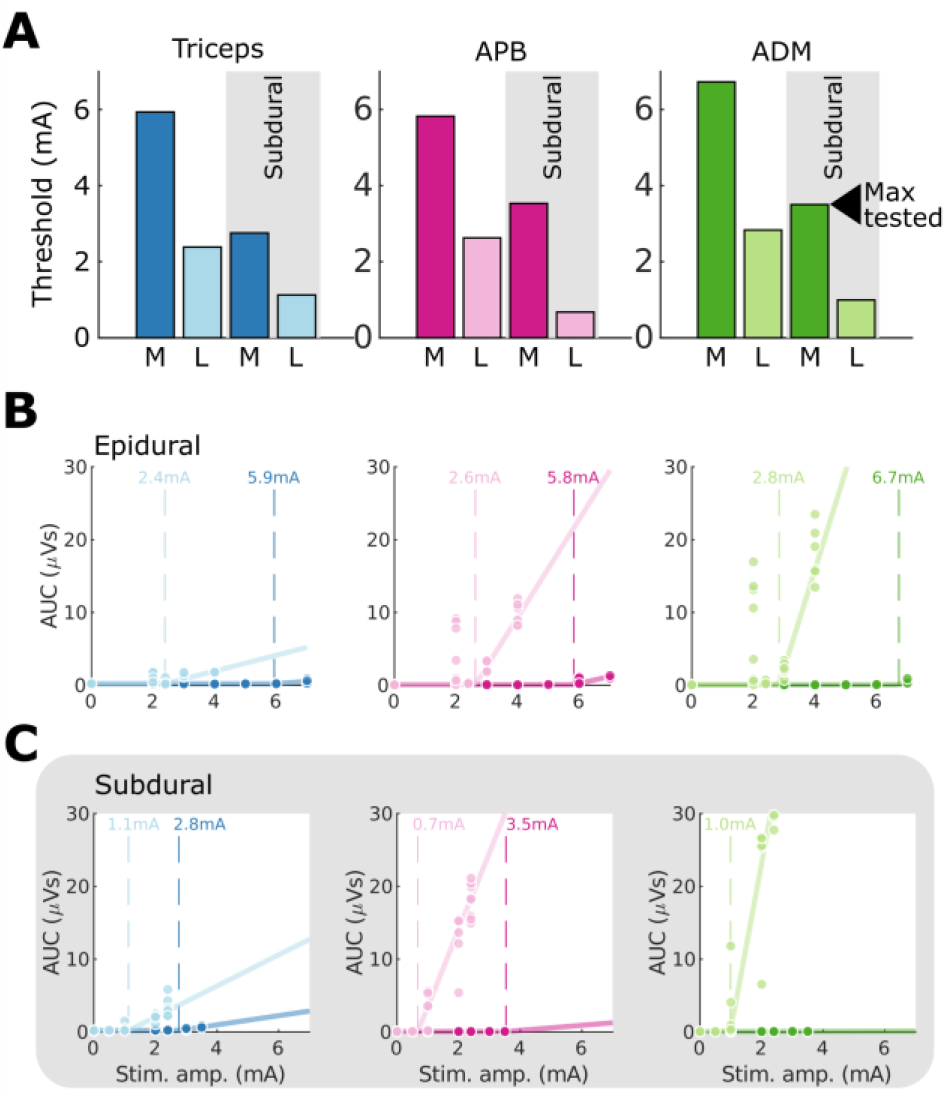
Comparison of subdural and epidural stimulation intensity thresholds at midline and laterally. **(A)** Summary of thresholds for midline (M, dark colors) and lateral (L, light colors) epidural and subdural stimulation. Lateral subdural stimulation is most effective, followed by lateral epidural stimulation, midline subdural stimulation, and midline epidural stimulation across the three activated muscles. **(B)** The threshold of muscle activation for epidural stimulation was determined by fitting functions to sections of data where multiple stimulation intensities were used. Circles indicate individual data points, solid lines show fitted functions, and dashed lines show estimates of threshold. **(C)** As for (B), with data from the corresponding subdural stimulation location.

#### 2.5.4 Comparison of responses at different cervical segments

All MEPs where the fixed intensity stimulation was used within a single participant were included in this analysis. The average AUC across participants was computed by using the geometric mean (Fig. 8A). The similarity in MEPs across segments was measured using the Spearman correlation coefficient, calculated between all segments across muscles for each participant (see Fig. 8B1-2) and then averaged across participants (see Fig. 8B3). To demonstrate similarity across segments, hierarchical agglomerative clustering was applied (38). This procedure recursively calculates the correlation between conditions and merges them to create a correlation between clusters which is displayed as a dendrogram. The similarity in MEPs across segments was quantified using the Spearman correlation coefficient, calculated between all segments across muscles for each participant and then averaged across participants. The merging procedure is based on the linkage metric which was conservatively set to the minimum distance (1 - correlation). We set the cluster cutoff at 70% of the maximum distance.

**Figure 8.**
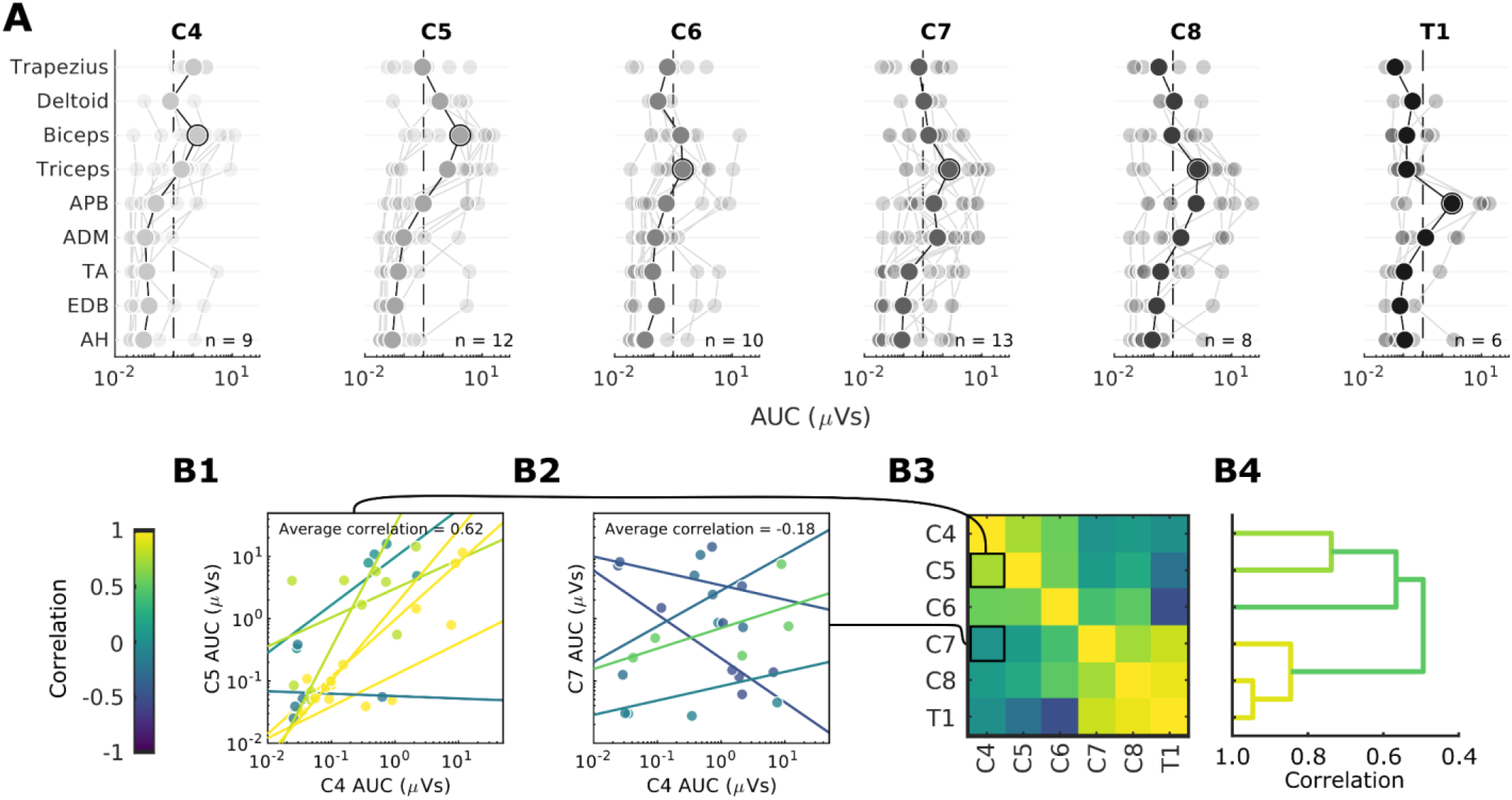
Rostral-caudal distribution of MEPs. **(A)** MEPs for each of the recorded muscles with stimulation are shown for lateral stimulation at segments C4-T1. The data are plotted on a log scale due to the exponential nature of MEPs to show their relative size. The average is shown in bold. Faint lines represent individual participant data (n = 25 total, with the number of participants represented shown within each panel). Dashed line (0.33μVs) indicates a threshold for presence of MEPs (see Methods). Equivalent figure with T2 signal change segments excluded is shown in Figure S1. **(B)** Within-participant similarity (correlation) of muscle responses at different segments, averaged across participants. (B1-2) Example AUC data correlations when comparing C4 to C5 (B1) or to a more distant C7 (B2). Nearby segments are highly correlated for the majority of participants (colors represent different participants), while more distant segments are not. (B3) Muscle activation appears to form a distinct cluster formed from lower cervical segments (C7, C8, T1). (B4) Dendrogram constructed from hierarchical clustering of (B3). Individual pairs of correlations are merged into clusters, and the maximum correlation between the entries in these clusters is represented by the width of the merge. Colors represent the maximum correlation within distinct clusters (see Methods).

#### 2.5.5 Midline-lateral versus rostral-caudal and anterior-posterior comparison of waveform shape and latency

To test the hypothesis that midline and lateral stimulation activate largely overlapping circuitry, an analysis of midline and lateral waveform shape (39) and onset latency was performed. Stimulation intensity impacts MEP size which is known to impact MEP shape (40). Consequently, it is important to match size across conditions before calculating similarity. For muscles of the arm and hand in all participants at the most caudal stimulated level, the fixed intensity MEP size at midline was matched to the MEP size in the lateral recruitment curve (a match was defined as an AUC difference less than 0.5μVs). If a match was found, two computations were performed to determine the overall similarity of the waveforms, independent of their size. First, the onset latency of these waveforms was computed based on the averages at each site (first deflection greater than 10μV). Second, the correlation of these waveforms was computed; this was done at multiple delays (from −12.5ms to 12.5ms) in order to determine similarity independent of differences in onset latency. The maximum correlation across these delays was then used to represent similarity.

In contrast to midline-lateral stimulation, we expected stimulation across different rostral-caudal segments to activate different circuits and consequently produce larger differences in onset latencies and waveform shapes. In order to calculate these metrics across segments, the same method developed to compare midline-lateral metrics was used to compare a fixed intensity MEP size at a rostral level to a size matched MEP at a more caudal level, where a recruitment curve was recorded. Waveform similarities and differences in onset latencies were compared between the midline-lateral condition and the rostral-caudal condition pooled across the segments. Due to the relatively low number of matches in MEP size, individual muscles were treated as independent from each other for this analysis (as indicated by the insets of Fig. 9A).

**Figure 9.**
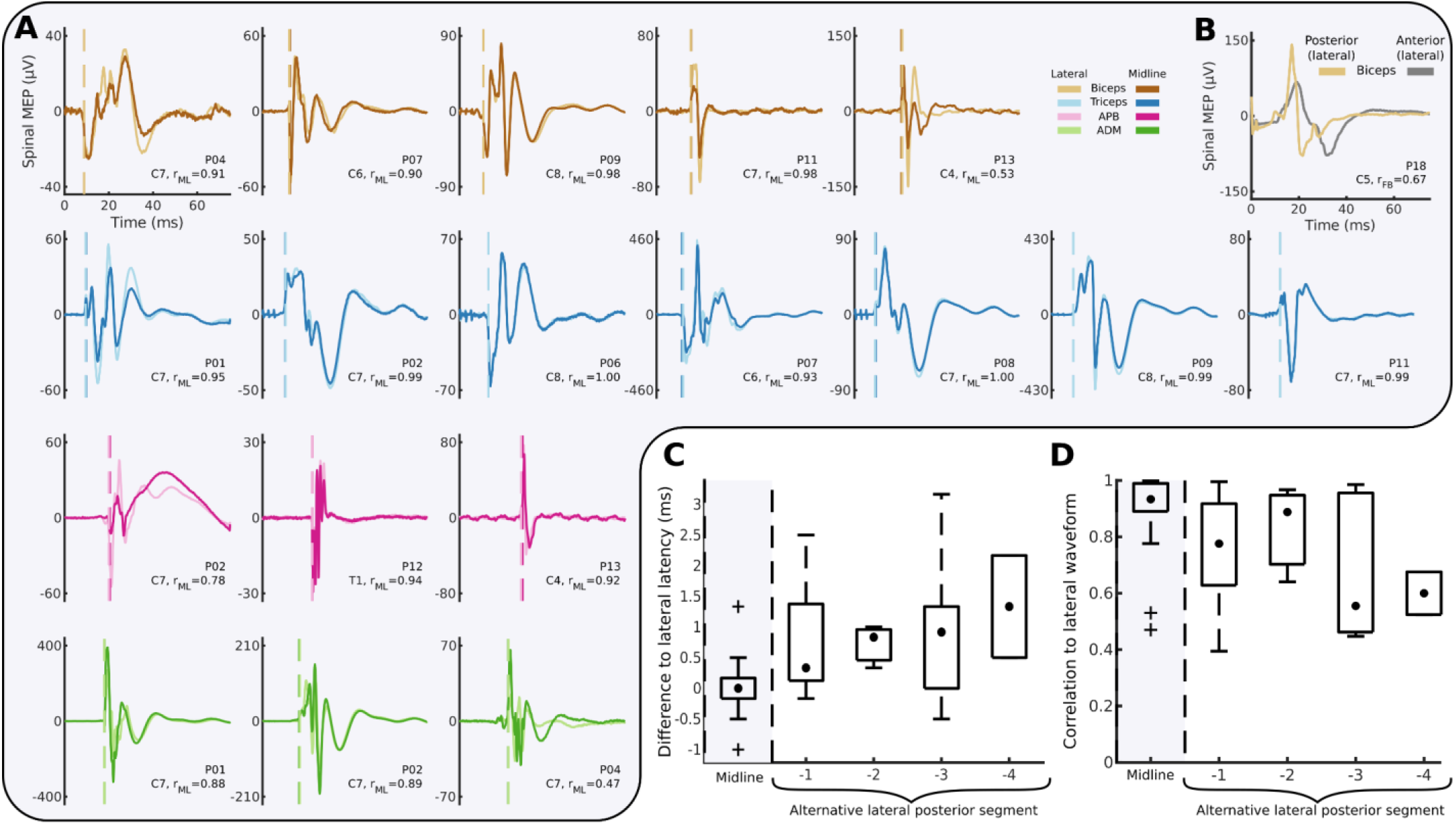
Comparison of midline-lateral, rostral-caudal and posterior-anterior waveform shapes. **(A)** Averaged midline (dark colors) and lateral (light colors) MEPs matched for size are highly similar. *r*_ML_ indicates the correlation between the two waveforms after adjusting for potential differences in onset latencies. **(B)** Waveform shapes for posterior-anterior comparison for a single participant in the biceps muscle appear highly distinct from each other. **(C)** Box plot of onset latencies for midline and lateral MEPs as in (A) (gray background), compared to onset latencies between lateral MEPs and MEPs at alternative lateral segments (white background). Midline-lateral latency differences are centered around 0ms, while latency differences increase with more distant segments. **(D)** Box plot of the averaged midline and lateral MEPs as in (A) (gray background), compared to correlations between lateral MEPs and MEPs at alternative lateral segments (white background). Midline-lateral correlations are higher than neighboring rostral-caudal correlations which appear to decrease at more distant segments. Hinges represent first and third quartile and whiskers span the range of the data not considered outliers (defined as q_3_ + 1.5 × (q_3_ – q_1_) or less than q_1_ – 1.5 × (q_3_ – q_1_)).

Finally, we tested the difference in responses from anterior and posterior spinal cord stimulation. This was done for two reasons: to determine whether responses are mediated by different pathways and to show that differences in responses would be detected by the waveform analysis. We expected that MEPs generated from lateral posterior stimulation would produce large differences in waveform shape when compared to anterior stimulation. This analysis was possible in a single participant where anterior and posterior segments were exposed and the dorsal root entry zone and corresponding ventral root location could be targeted. To determine whether waveform shape was more or less different in the anterior-posterior condition compared to the midline-lateral condition, the anterior-posterior similarity metric was ranked relative to the midline-lateral metrics and reported as a percentile. The latency difference was not calculated for this condition because stimulation artifacts caused baseline shifts that made the onset time estimation unreliable.

#### 2.5.6 Estimating the change in MEP produced by moving electrodes midline to lateral versus between segments

Comparisons between movement of electrodes along both the midline-lateral and rostro-caudal axes was possible in 9 participants. The segment of maximum AUC caused by lateral stimulation was first identified for each muscle group; subsequently the differences between MEP sizes at this and neighboring segment(s) were calculated. If two neighboring segments were present, the two differences were calculated and then averaged. Similarly, the difference between the lateral MEP and midline MEP sizes at the same segment was calculated for each muscle. The differences in MEPs were then normalized by distance by dividing MEP values by a generic inter-root length for the rostro-caudal axis and half the transverse cord diameter for the midline versus lateral placement. Distance estimates were derived from a human cadaveric study (41), in which inter-root distances in the cervical spinal cord (average 12.5mm) and midline to lateral distance (average 6.7mm) were measured. Each participant had the MEP sizes across muscles averaged; a Wilcoxon signed-rank test was applied to determine the relative change along each axis.

#### 2.5.7 Estimation of threshold and slope

Recruitment curves were collected at only one segment per participant because of time constraints. Recruitment curves were fit to enable comparison of slope and threshold across multiple segments. Specifically, the stimulation intensity and AUC relationship was modeled as a softplus function: 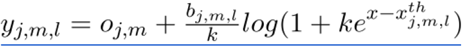, where <u>*y_j,m,l_*</u> represents the AUC for a given participant <u>(*j*)</u> muscle <u>(*m*)</u> and electrode location <u>(*l*)</u> for a given stimulation intensity <u>*x*</u> Symbols <u>*o*</u>,<u>*d*</u>, and <u>*x^th^*</u> represent the recruitment curve offset, slope and threshold respectively. The parameter *<u>k</u>* was set to 20 to approximate a linear rectified function while maintaining numerical stability. A fit was performed with a generalized pattern search algorithm (see Supplemental Methods) for every muscle and participant simultaneously for multiple locations on the spinal cord (due to the shared offset). A shared offset was used because a subset of data did not test a stimulus intensity below MEP threshold. In order to compare thresholds across the midline and lateral conditions across participants, a Wilcoxon signed-rank test was used on the difference between midline and lateral estimates averaged across segments and muscles. The same procedure was used for a comparison of the slope estimates.

## 3 Results

### 3.1 Participant recruitment and characteristics

Participants (n = 25, twelve male and thirteen female) underwent surgery that exposed the cervical enlargement for electrical stimulation; their demographics and clinical findings are summarized in Table 1. Mean age of study participants was 65.5 years (range 39 to 83). Twenty-two patients had chronic symptoms, and three patients had subacute symptoms. Twenty-one patients had clinical signs of myelopathy or myeloradiculopathy (22 had T2 signal change on MRI, indicating myelopathy); the remaining patients had radiculopathy alone.

### 3.2 Location of stimulating electrodes

The location of the stimulating electrodes was determined two ways. First, electrode location based on anatomical landmarks (Fig. 2A1) was identified in an intraoperative CT (Fig. 2A2) and then coregistered (Fig. 2A3) into MRI space in one participant. The location of the electrode was confirmed to be over the C7 dorsal root entry (Fig. 2A4), where it had been placed intraoperatively.

As a secondary confirmation of the electrode placement strategy, an additional experiment was performed in a patient undergoing laminectomy and dural opening for resection of intradural meningioma. Electrodes were placed relative to anatomical landmarks as for epidural stimulation, and once the dura was removed confirmation was made that the location matched the dorsal root entry zone (Fig. 2B).

### 3.3 Lateral stimulation is more effective than midline stimulation

Comparison of spinal motor evoked potentials (MEPs) to stimulation at midline and lateral locations was performed in 14 participants. We confirmed that placing the electrodes in the lateral position located them over the dorsal root entry zone, where afferent axons enter the spinal cord (Fig. 2). Fig. 3 shows MEPs in response to midline (dark) or lateral (light) stimulation, using an intensity set to 120% of the threshold for midline MEPs. Lateral stimulation of the cervical spinal cord generated larger MEPs than midline stimulation across participants (Fig. 3A; mean midline AUC = 1.14±0.82μVs, lateral AUC = 4.71±2.28μVs, Wilcoxon signed-rank test, *p* = 0.002, n = 14). These changes correspond to a median increase of 258% in the MEP when stimulating laterally versus at midline. The subplots in Fig. 3B show raw MEPs of the 14 participants. The upper insets show violin plots of the lateral and midline MEPs for each individual, with asterisks indicating significant differences *(p* < 0.05, Bonferroni corrected Wilcoxon rank sum test). Ten of 14 participants showed significantly increased MEPs with lateral stimulation compared with midline.

The greater effectiveness of lateral compared to midline stimulation was not unique to a specific muscle or any particular segment. Fig. 4 shows each of the six arm muscles in which MEPs were recorded and the six spinal segments that were stimulated. Each dot represents the lateral (y-axis) and midline (x-axis) responses to stimulation at the various segments in each of the participants. All responses above the x = y line represent larger lateral than midline MEPs. On average, across participants, segments and muscles, 91.5% of MEPs were larger when the spinal cord was stimulated laterally.

The increased effectiveness of lateral stimulation in the cervical cord was also observed in the leg muscles. Fig. 5A shows the summary of responses for midline (dark) or lateral (light) stimulation. In the 6 out of 14 participants who had tibialis anterior (TA) MEPs in response to cervical stimulation, lateral cervical stimulation generated larger leg MEPs than midline stimulation (AUC = 0.11±0.10μVs, lateral AUC = 2.43±1.54μVs, Wilcoxon signed-rank test, *p* = 0.031, n = 6). The subplots in Fig. 5 show raw MEPs of the 6 participants. The upper insets show a violin plot of the lateral and midline MEPs for that individual (*p* < 0.05, Bonferroni corrected Wilcoxon rank sum test). All of the 6 participants showed significantly increased MEPs with lateral stimulation compared with midline.

### 3.4 Lateral MEPs are larger due to lower thresholds and steeper recruitment curves

The increased effectiveness of lateral stimulation may be driven by a reduction in the threshold for recruitment, an increase in the rate of change of MEP size with stimulation intensity (slope of the recruitment), or both. Recruitment curves were fitted in order to estimate these parameters directly from data where multiple stimulation intensities were tested at the same segment (e.g. Fig. 1C, C7). Examples of the recruitment curve threshold and slope estimation procedure for the triceps muscle of two participants are shown in Fig. 6A for midline and lateral stimulation. Across participants, average threshold increased 107% for midline stimulation over lateral stimulation, indicating higher stimulus intensity was needed to evoke an MEP *(p* = 2×10^-4^, Wilcoxon signed-rank test, n = 13; Fig. 6B). The rate of increase of MEP size with stimulation intensity was also influenced by the site of stimulation, with a median increase in the slope of 34% for lateral stimulation compared to midline stimulation *(p* = 0.010, Wilcoxon signed-rank test, n = 13; Fig. 6C). Thus, lateral stimulation was more effective than midline stimulation both because of a lower threshold and because of an increased slope of the recruitment curve.

### 3.5 Lateral stimulation efficacy is most pronounced when applied under the dura

Stimulation was applied midline and laterally below the dura at the T1 root level, and at matched epidural locations. Across the three muscles activated (Fig. 7A-B), the lowest intensity stimulation was found for stimulation of the dorsal root entry zone below the surface of the dura, followed by lateral epidural, midline subdural and lastly midline epidural stimulation locations.

### 3.6 Cervical dorsal stimulation evokes MEPs in muscles innervated at that segment but also muscles innervated at adjacent and remote segments

While we predicted a sharp difference in responses between midline and lateral stimulation, we expected smaller differences with movement between cervical segments along the rostro-caudal axis (Fig. 1B). MEPs were recorded in arm and leg muscles after a fixed intensity stimulation at 6 different spinal segments (C4-T1). Fig. 8A shows MEP values for each muscle with stimulation at the segments available for a particular participant. Individual participants are represented by faint lines, and the group average for each muscle by bold lines. The dashed lines indicate a threshold for the presence of MEPs. The largest MEPs were observed in the muscles innervated at the stimulated segment. For example, biceps activation was dominant at C4 and C5, triceps at C6, C7 and C8, and APB at T1. Triceps MEPs were present at all segments except for T1. Additionally, as shown in Fig. 5, some participants had MEPs in the legs, although those were always smaller.

The patterns of responses are likely to reveal how similar each segment is to the other segments. To determine this, we calculated the correlation between segments of MEPs from all muscles. Fig. 8B1 or B2 show the results of all the individual participants that had stimulation at the two segments indicated on the axes. Each dot corresponds to the MEP for an individual muscle, and all the arm and hand muscles for an individual are used to create a linear correlation represented by a line. By averaging these correlations across participants (Fig. 8B3), we were able to examine the similarity of muscle responses across segments without the confound of across-participant variability. MEPs from C7-T1 largely fell into a cluster more distinct from rostral levels. We used a clustering algorithm to show these relationships; the correlation within the C7-T1 cluster averaged 0.85, while the correlation within the C4-C6 cluster averaged 0.57. The correlation between the rostral and caudal clusters was 0.5. Thus, the caudal cluster is strongly related internally, with much lower correlations to other levels.

Our goal was to understand the organization of the intact cervical spinal cord, but the study was performed in people with myelopathy. To control for the effect of injury to the spinal cord, we performed lateral stimulation on the less affected side. In addition, we performed an analysis similar to Fig. 8 but excluding responses in all segments where T2 signal change was present. This did not change the pattern of responses significantly (average correlation between MEP size with and without T2 signal change segments excluded *r* = 0.89, Supplemental Fig. S1). This high correlation suggests that the patterns of muscle activation from cervical spinal cord stimulation were driven largely by the activation of intrinsic circuits that were spared from the effects of myelopathy.

### 3.7 Similar latencies and waveforms from midline and lateral stimulation matched for size

We expect that both midline and lateral stimulation recruit large-diameter afferent axons as they enter the spinal cord. While lateral stimulation was far more effective at producing MEPs, we expected that the latency and waveform of the resultant MEPs from stimulation in each location would be similar when matched for the size of the MEP (see Methods). We examined this by comparing variation in latencies and waveform shape correlations along the midline-lateral (Fig. 9A) and rostral-caudal axes. For the single participant that underwent anterior and posterior surgery, the waveform shape of the biceps MEPs from the two stimulation aspects appear highly distinct (Fig. 9B), and this is reflected in a low correlation (*r* = 0.67, 11^th^ percentile of the posterior midline and lateral MEP correlations). The difference in onset latencies between midline and lateral responses was nearly identical (0.04 ± 0.22ms), and this was not different from 0ms *(p* = 0.655, n = 18). In contrast, the latency from lateral responses at various levels was different (Fig. 9C), with longer latencies from progressively more distant segments; pooled across segments, the latency was different from the midline-lateral condition (0.83 ± 0.22ms, n = 24, *p* = 0.002). We also tested the similarity in waveforms between conditions. MEP waveforms were very similar between midline and lateral stimulation (*r* = 0.89 ± 0.04, Fig. 9D). The similarity became less across more distant segments (Fig. 9D); waveforms were more similar for the midline-lateral correlation than across segments (pooled *r* = 0.77 ± 0.05, n = 24, *p* = 0.029).

### 3.8 Map of MEPs produced by epidural stimulation of the cervical spinal cord

Fig. 10A summarizes the muscles activated by stimulation at each cervical segment. Circle area corresponds to the percent of total MEP size contributed by specific muscles (designated by color), and the darkness of the circle corresponds to the number of participants. Only MEPs that were on average greater than 0.33μVs are shown, highlighting the dominant muscles at each segment. While all lateral stimulation sites met this threshold, the only midline response above this threshold was the triceps MEP at C7 and C8. This map demonstrates the activation of specific muscles at each segment of the cervical spinal cord.

**Figure 10.**
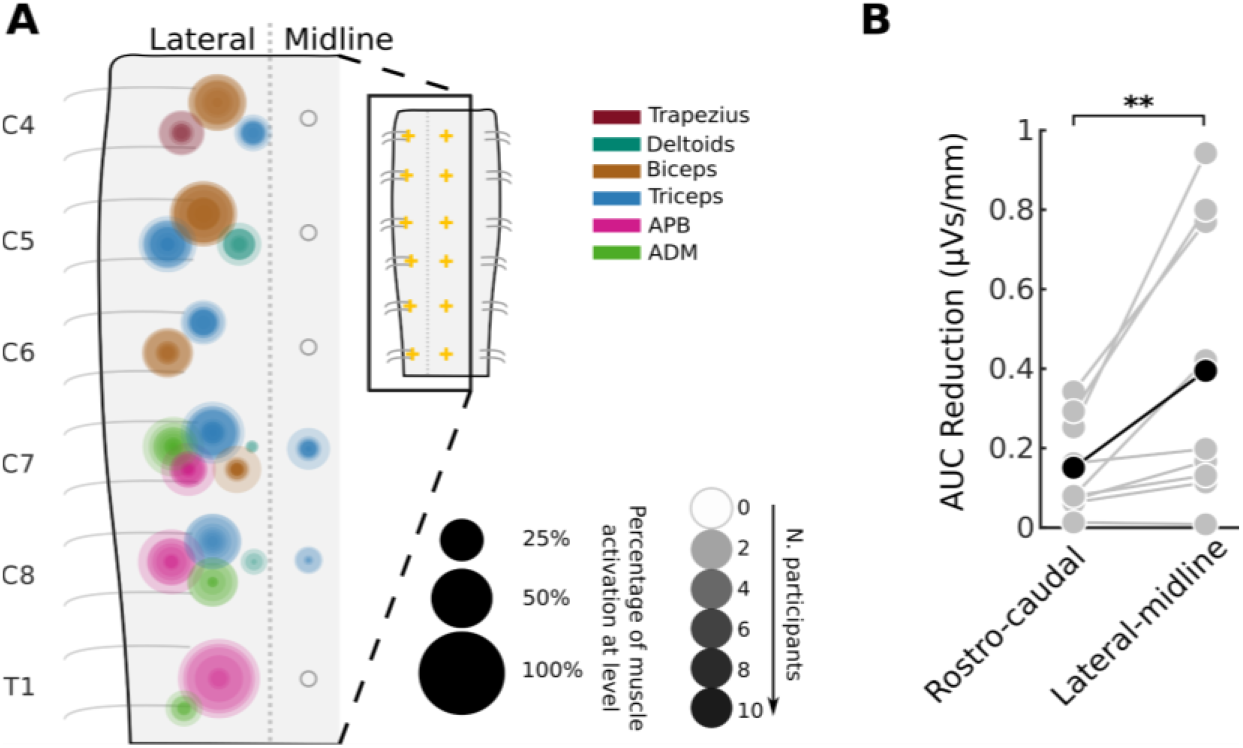
An actionable map for epidural stimulation of the cervical spinal cord. **(A)** Area of overlaid circles represents the rectified area under the curve (AUC) of individual participants. Activation is only drawn when the across-participant average AUC is greater than 0.33μVs (see methods). Empty circles represent no muscle activation reached this threshold. (Data shown here is a summarized representation of data shown in Fig. 8A). Inset: Crosses mark locations of stimulation. **(B)** Greater change in AUC in the lateral-midline axis as compared to the rostro-caudal axis. Individuals are shown as gray dots with lines connecting their values, and the average of the participants in black.

Differences between MEP size were larger between midline and lateral stimulation within segments than between segments. To quantify this, we determined change in MEPs as a function of distance – how much does the MEP change with each millimeter that the stimulating electrode was moved in each direction. The average MEP size was reduced at a rate of 0.39±0.24μVs/mm when moving from lateral to midline. On the other hand, MEP size from stimulation of neighboring segments was reduced at a lower rate of 0.15±0.08μVs/mm. The rate of change was more than double for lateral-midline electrode position compared with rostro-caudal position, and this difference was highly significant (*p* = 0.008, n = 9).

## 4 Discussion

In addition to the empiric map, these results demonstrate the underlying organization of the human cervical spinal cord. The largest changes in stimulus response were observed between midline and lateral locations which is likely due to the greater efficacy of recruitment when the electrodes were placed over the dorsal root entry zone. Consistent with this, the relative efficacy of lateral stimulation was maintained when stimulating below the dura over the dorsal root entry zone. Movement of stimulating electrodes along the rostro-caudal axis resulted in more subtle differences in motor responses, suggesting the presence of strong overlap with adjoining segments. Clustering of these responses was particularly strong within lower cervical segments (C7, C8, T1), with more diversity at other levels. Finally, we observed MEPs arising from distant segments of the cervical spinal cord and from leg muscles, indicating potential activation of propriospinal connections (16, 42). Thus, the summary map of responses (Fig. 8A) provides a guide to target muscles via dorsal cervical spinal cord stimulation but also an organizational map of neural connections that can be recruited with this technique.

The observed differences in MEP size between midline and lateral stimulation and between neighboring segments have support in the literature. However, our work provides novel electrophysiological evidence that midline and lateral stimulation predominantly activates the same circuitry, while stimulation at more distant lateral rostral-caudal sites incorporates additionally more diverse circuits. The significantly different MEP morphology in response to anterior stimulation suggests that the MEPs generated in response to dorsal stimulation are indeed mediated through the dorsal afferent pathways rather than through direct activation of the motor neuron via spread to the ventral horn. Unexpected spread of responses beyond the stimulated segment has been reported (31), even with intraspinal cervical stimulation (36, 43). Activation of leg muscles from stimulation of the cervical cord has also been found (31–33, 44). Consistent with these findings we observed long-range activation, for example triceps activation with C4 stimulation and leg activation with cervical stimulation. Such activation cannot be explained by known locations of motor pools and intermingling of motoneuron cell bodies (26, 27) and suggests the presence of long distance connections within the cervical enlargement (29, 45, 46) and between the cervical and lumbar enlargements (16, 47–50).

Clustering of lower cervical responses may reflect the intrinsic organization of the spinal cord in humans. In the monkey cervical cord, Greiner et al. demonstrated separation of biceps and triceps by comparing MEPs from stimulation at the C6-C8 segments. However, consistent with human data also described by Greiner et al., our results suggest that this distinction may be less pronounced in bipedal humans compared with quadrupedal macaques due to the widespread activation of the triceps muscle. Rather, our results suggest a division between the muscles of the hand and muscles of the arm. This organization could be engaged to foster adaptive patterns of arm and hand muscle activation.

The patterns of muscle activation from dorsal spinal cord stimulation can be provisionally inferred through known spinal circuits. Spread between adjacent segments (for example, strong activation of biceps from stimulation at C4) may be mediated by the spread of afferent connections into the spinal cord or the spread of longitudinally extensive motor pools out of the spinal cord. Some observed responses may be partially driven by non-biological spread of the electric field across levels (21), but the presence of leg muscle MEPs provides strong evidence that spinal cord pathways represent a critical mechanism. Additionally, we observe a much larger change in MEP size when electrodes are moved in the midline-lateral orientation than when they are moved in the rostral-caudal orientation, suggestive of the limitations of current spread.

While the operating room provides a unique experimental setting, it may also limit the generalizability of our findings. Participants had myelopathy and/or radiculopathy, and these pathologies may affect axonal conduction, neuronal signaling, or synaptic connectivity. We did, however, perform a sensitivity analysis for the effect of cervical injury by excluding the segments with T2 hyperintensity, indicative of myelomalacia; excluding these levels did not change the overall structure of the activation map. MEP size may also have been affected by general anesthesia; however, all participants were maintained on total intravenous anesthesia, consistent with standard operative and intraoperative neuromonitoring conditions; in all our experiments we observed clear, reproducible responses at multiple tested segments. Importantly, anesthetic conditions had reached a steady state during recording, and no changes were made during the experiment. Because participants were tested in an immobilized position on the operating table, we were not able to address how epidural stimulation can be used to generate movement (23, 44, 51) or how MEPs interact with body position (52–55). Normalization of MEPs by their maximal value is indicative of the proportion of recruited motoneurons and less impacted by sources of variability such as muscle size and subdermal electrode position. However, MEPs could not be consistently driven to saturation under intraoperative conditions, and data has consequently been presented without normalization which may have led to some over or under representation of specific muscles. Muscle selection is chosen based on the cervical nerve roots which are most at risk of iatrogenic injury due to surgical maneuvers. Spinal cord function can be adequately monitored with a minimal amount of upper and lower extremity recording sites, but multiple muscle groups are required to be nerve root specific. Specific muscle groups are chosen based on accepted myotome patterns and clinical review of surgical outcomes when compared to data changes. Because wrist muscles were omitted, the observed rostral-caudal clustering of muscle activation is likely to be only partially reflective of the clustering of the circuitry of the spinal cord. Stimulation technique or study population may also be critical for the specific form of the observed clustering. In future studies these limitations may be addressed in alternative experimental settings either non-invasively or with implanted leads.

The map of muscle responses to cervical epidural stimulation can facilitate better understanding of spinal circuits and help target interventions. Stimulation of the cervical spinal cord may reveal patterns of activation, whether mediated by motor primitives or complex movements. In addition to single or short trains of stimuli, longer trains and/or multisite stimulation may help to reveal these motor programs. Multiple sites of stimulation may be needed to activate the desired circuit activation or movement. This map may be used to guide further experiments to elucidate optimal sites and stimulation patterns for activating movement and strengthening damaged spinal circuits.

## 5 Data availability

The data that support the findings of this study are available from the corresponding authors, upon reasonable request. Supplemental material is available at DOI: https://doi.org/10.6084/m9.figshare.19891966.

## Abbreviations

mJOA: Modified Japanese Orthopaedic Association;
MEP: Motor Evoked Potential;
TA: Tibialis Anterior;
EDB: Extensor Digitorum Brevis;
AH: Abductor Hallucis;
AUC: Area Under the Curve;
SCI: Spinal Cord Injury;
SEM: Standard Error of the Mean.

## Acknowledgements

Sources of financial support: This study was funded by the National Institutes of Health (1R01NS124224); and the Travis Roy Foundation Boston, MA (Investigator Initiated).

We thank neurologists P. Kent, H. Choi, M. Bell (The Och Spine Hospital At New York Presbyterian Hospital) and intraoperative monitoring technologists N. Patel, Z. Moheet (Weill Cornell Medicine), Joe Elliott, Brian Demboski, Kelley Wichman, Susannah Storms, Meghan Mullaney, Evance Desriviere (The Och Spine Hospital At New York Presbyterian Hospital) for monitoring patient safety during the experiments, as well as help with running the experiments. We also thank S. Oh (Columbia University), E. Wong (Brainlab AG) and Brainlab AG for help with image processing and M. Vulapalli, C. Mykolajtchuk, M. Michael (Weill Cornell Medicine) for help in administrative matters.

