## Supplementary material for "Intraoperative electrical stimulation of the human dorsal spinal cord reveals a map of arm and hand muscle responses": Table 1-1 (from another version of this preprint: version 1, 2022-01-31)

| **Table 1-1. Experiment parameters** | | | | | | |
|---|---|---|---|---|---|---|
| | Most responsive muscle* | Conditions tested | Figures | Stimulation waveform | Pulsewidth | Inter-stimulus interval |
| P01 | ADM | **C7M, C7L** | 3-8 | biphasic | 250µs | 2ms |
| P02 | Triceps | **C7M, C7L, C6M, C6L, C5M, C4M** | 3-8 | biphasic | 250µs | 2ms |
| P03 | Triceps | **C7M, C7L, C6M, C5M, C4M, C4L, C4L** | 3-8 | biphasic | 250µs | 2ms |
| P04 | Triceps | C7M, C7L, C4L | 3-5, 7-8 | biphasic | 250µs | 2ms |
| P05 | Triceps | **C8M(U), C8L(U), C7M(U), C7L(U), C6M, C6L, C5M, C5L** | 3-8 | biphasic | 250µs | 2ms |
| P06 | Triceps | **C8M, C8L, C7M, C6M, C5M, C5L, C4M** | 3-8 | biphasic | 250µs | 2ms |
| P07 | Triceps | C8M, C8L, **C7L, C6M, C6L**, C5M, **C5L** | 3-8 | biphasic | 250µs | 2ms |
| P08 | Triceps | **C7M, C7L,** C6M, C6L, C5M, C5L, C4M, C4R | 1, 3-8 | biphasic | 250µs | 2ms |
| P09 | Triceps | **C8M, C8L,** C7M, C7L, C6M, C6L, C5M, C5L, C4M, C4L | 3-8 | biphasic | 250µs | 2ms |
| P10 | Triceps | **C8M(U)**, **C7M, C7L**, C6M, C6L, C5M, C5L, C4M, C4L | 3-8 | biphasic | 250µs | 2ms |
| P11 | Triceps | **C8M(U)**, **C7M, C7L,** C6M, C6L, C5M, C5L, C4M, C4L | 3-8 | biphasic | 250µs | 2ms |
| P12 | APB | **T1M, T1L, T1L,** C8M, C8L, C7M, C7L | 3-8 | biphasic | 250µs | 2ms |
| P13 | Biceps | **C4M, C4L**, C4L, **C4L* C4L*** | 3-8 | biphasic | 250µs | 2ms |
| P14 | APB | **C8M, C8L, C8L, C7L, C6L, C5L, C4L** | 3-8 | monophasic | 500µs | 2ms |
| P15 | Trapezius | C5M, C5L, C5L*, C4M(U), C4L | 8 | biphasic | 250µs | - |
| P16 | APB | C8L, C5L, T1L | 8 | biphasic | 250µs | - |
| P17 | Triceps | C6L*, **C5L, C5L*, C4R** | 8 | monophasic | 300µs | - |
| P18 | Biceps | **T1L***, T1L*, **C6L, C5L** | 8 | monophasic | 300µs | - |

Conditions tested: Plaintext entry indicates segment (e.g. C7) and midline or lateral location (M or L) with 3 pulse stimulation and handheld ball electrode. Bold indicates multiple tested stimulation intensities, * indicates catheter electrode, underlined indicates single pulse. (U) indicates level was non-responsive (AUC < 0.33µVs) at tested stimulation intensities (up to 10mA max). Inter-stimulus interval is applicable to all 3 pulse data. P01-P10, P15 were recorded with a Cadwell Elite/Pro (Cadwell Inc.), P11-P13, P16 were recorded with a Cadwell IOMAX (Cadwell Inc.), P14, P17, P18 were recorded with an XLTEK Protektor32 (Natus Medical Inc).
