## Supplementary material for "Intraoperative electrical stimulation of the human dorsal spinal cord reveals a map of arm and hand muscle responses": Figure 7-1 (from another version of this preprint: version 1, 2022-01-31)

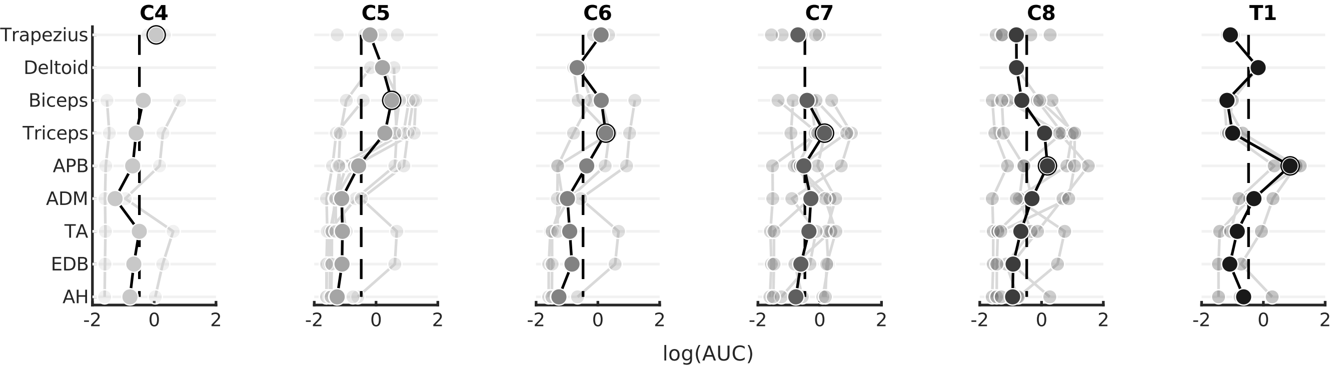


**Figure 7-1. Rostral-caudal distribution of MEPs excluding MEPs at segments where T2-weighted signal change is present.** Average distribution of the rectified area under the curve (AUC) of the spinal motor evoked potential (MEP) across muscles at different cervical segments (solid line). Stimulation is applied laterally over the dorsal root entry zone. Transparent lines in the background correspond to individual participant data (n = 18). Across-segment average correlation of averaged responses with data that does not exclude segments with T2-weighted signal change is high (r = 0.91).
